# Cytoplasmic polyadenylation protein-3 is expressed in adult mouse cholinergic spinal neurons, including motoneurons

**DOI:** 10.64898/2026.11.23.767601

**Authors:** Pei San Lew, Prabhisha Silwal, Shenani Basnayake, Felicia Sinclair, Sharma Bhavya, Xiaoyu Chen, Katinka Stecina

## Abstract

Activity-dependent plasticity occurs in spinal networks, but mechanisms underlying the retention or modification of spinal synapses are less well understood than those in the brain. The cytoplasmic polyadenylation protein 3, or CPEB3, is one of four variants discovered in mammals that change in response to motor activity in hippocampal neurons. Here, we examined whether CPEB3 is expressed in spinal neurons and whether changes in CPEB3 following locomotor activity can be detected. Low thoracic and lumbar spinal cord segments were examined by immunoblotting of homogenized spinal tissue (n=6 mice, 3 female). We compared samples from mice subjected to 2 hours of treadmill walking with those from mice that did not walk by western blotting. There was significantly more CPEB3, by 20.7 +/- 15%, in soluble fractions obtained from the thoracic spinal cord of male mice that performed walking compared to control mice. In male mice, STAT5b increased by 6.8 +/- 0.1% and by 15.7 +/- 0.1% in thoracic and lumbar soluble fractions, respectively. In samples from female mice, variability was greater, and walking did not produce significant changes. Immunofluorescence microscopy in another cohort of mice (n=6) showed that 88 +/- 0.1% of NeuN-positive cells in the ventral horn were also positive for CPEB3. Moreover, CPEB3 expression was evident in ventral cholinergic neurons, including motoneurons. These findings establish that CPEB3 is present in cholinergic spinal neurons of the ventral horn, and this warrants further exploration in motor-activity-associated spinal plasticity.

## Introduction

The ability of synapses to undergo morphological and biochemical changes in response to stimulation, which is referred to as synaptic plasticity, is a foundational mechanism underlying learning and memory. Maintaining long-term memory requires *de novo* protein synthesis and depends on translational regulation, as reviewed by e.g. (Citri & Malenka, 2008). One critical pathway of post-transcriptional control is cytoplasmic polyadenylation, a process that activates the translation of dormant mRNAs (Hake & Richter, 1994). This mechanism is widely recognized as a key component of neural plasticity in vertebrates (Hicks et al., 2025). Among the primary mediators of this process is the cytoplasmic polyadenylation element-binding (CPEB) protein family, which consists of several nucleus-cytoplasm shuttling RNA-binding proteins. CPEBs bind to the consensus 3-prime UTR cytoplasmic polyadenylation element (CPE; UUUUUAU) of target mRNAs to regulate polyadenylation-induced translation during cell division, cellular senescence, and synaptic plasticity (Barnard et al., 2004; Hake & Richter, 1994; Richter, 2007). In mice, four distinct CPEB proteins (CPEB1–4) are expressed. Of these, CPEB3 acts as a sequence-specific repressor of target neural mRNAs in the basal state but promotes translation following neural activity (Fioriti et al., 2015; Ford et al., 2019; Richter, 2007). Due to its interactions with the cytoskeleton, CPEB3 acts as a functional prion-like protein essential for maintaining long-term memory storage in the murine hippocampus (Fioriti et al., 2015; Stephan et al., 2015; Theis et al., 2003) and CPEB3 has been implicated in human episodic memory (Vogler et al., 2009). Following NMDA receptor stimulation, CPEB3 localizes to the nucleus of hippocampal neurons (Chao et al., 2012) and its synaptic levels increase within 30 minutes, dynamically modulating synaptic proteins such as GluR2 (Huang et al., 2006). Emerging evidence also highlights CPEB3’s unconventional role within the neural nucleus. Upon nuclear translocation, CPEB3 interacts directly with the transcription factor Signal Transducer and Activator of Transcription 5b (STAT5b) and this interaction serves as a pivotal regulatory switch for downstream signaling pathways, such as the STAT5b-linked transcription leading to drive AMPA receptor expression (Peng et al., 2010).

While activity-dependent plasticity and memory formation are well studied in cortical and hippocampal pathways, spinal neural circuits also undergo profound short- and long-term functional changes (Grau & Hook, 2006; Wolpaw, 2010) yet the mechanisms underpinning spinal cord memory are less understood than those of brain circuits (Guertin, 2008). Changes and interactions of protein cascades within spinal neurons following motor tasks remain far less explored than interactions in the brain. Motor control and motor memory are intrinsically tied to downstream activity in the spinal cord, particularly at the ultimate point of motor output: the spinal motoneuron. As the spinal cord executes automatic, learned, and skilled movements, new synaptic connections are induced in both animals (Grau et al., 2006; Grau & Hook, 2006) and humans (Vahdat et al., 2015) within spinal networks in response to motor activity. Locomotion serves as a classic paradigm to examine this activity-dependent plasticity within spinal networks (Grau et al., 2006). Neurons activated during different locomotor tasks can be mapped via immediate-early gene expression, such as *c-Fos*(Ahn et al., 2006; Dai et al., 2005; Dai et al., 2009), revealing diverse, active neural populations that include cholinergic neurons within the thoraco-lumbar segments of the cord (Noga et al., 2009, 2011).

Beyond the brain, CPEB3 has been reported in the murine retina, specifically in cholinergic amacrine cells during postnatal development (Wang & Cooper, 2009) and in the spinal dorsal root ganglia, where it modulates TRPV1 translation and nociceptive processing (Fong et al., 2016). To our knowledge, however, CPEB3 expression and dynamics have not yet been examined in relation to spinal motor networks. Therefore, this study aimed to investigate the expression and localization of CPEB3 in thoraco-lumbar neurons following a locomotor task. We examined whether CPEB3 is expressed within cholinergic murine spinal neurons and hypothesized that sustained locomotor activity triggers a relative increase in both CPEB3 and Stat5b in soluble (cytosolic) fractions collected from thoraco-lumbar spinal cord segments.

## Methods

The procedures and techniques conformed to ethical and scientific standards set by the Canadian Council on Animal Care and the University of Manitoba Central Animal Care Committee approved all protocols before the experiments began. In total, 18 mice were used; for western blotting 12 C57BL/6 mice (6 females, 19-22 g) and for tissue-sectioning and immunohistochemical identification of proteins 6 cfos-EGFP mice (4 females, 16-21 g) were used (Barth et al., 2004; Dai et al., 2005). All animals were from an in-house colony, group housed in 12-hr dark/light cycles with *ad libitum* access to food and water.

### Locomotor activity task

The C57BL/(n=12) were randomly assigned among cage mates to the locomotor or the control group. Mice assigned to the locomotor group were placed on a rodent treadmill, while the other mouse (control) remained in the home cage. Mice assigned to the locomotor group explored the new environment for 2-3 min before the treadmill speed was increased to 5 cm/s. Then the speed was adjusted to be between 8 and 15 cm/s. This adjustment was carefully monitored by the experimenter; to encourage a consistent walking pattern. All sessions consisted of 10-20 min bouts of walking alternating with 2-3 min breaks when treadmill belt speed was zero. A water bottle and small food pellets were in the front end of the treadmill, accessible to the animal. All locomotor mice were on the treadmill for a total of 120 min, and the distance walked is reported in Table 1.

**Table 1.** Description of mice and sample collected for western blotting. Sex and age (days) of mice used, distance walked on treadmill (m) over 120 minutes and the thoracic (T) and lumbar (L) sample’s length (l, mm) measured during harvesting and the time taken in minutes (t, min) from the start of the cardiac puncture in the perfusion procedure to freeze the sample, calculated with means and standard deviation (SD) for each group.

| Locomotor group |  |  |  |  |  |  |
|---|---|---|---|---|---|---|
| **Sex** | **Age (day)** | **walk (m)** | **T -l mm** | **T-t min** | **L-l mm** | **L-t min** |
| male | 50 | 735 | 7 | 19 | 7 | 13 |
| male | 51 | 700 | 6 | 17 | 6 | 12 |
| male | 67 | 857 | 6 | 21 | 7 | 20 |
| **mean** | **56** | **764** | **6** | **19** | **7** | **15** |
| **SD** | **10** | **82** | **1** | **2** | **1** | **4** |
| female | 146 | 713 | 6 | 20 | 7 | 16 |
| female | 78 | 772 | 7 | 17 | 5 | 11 |
| female | 98 | 671 | 6 | 18 | 7 | 21 |
| **mean** | **107** | **719** | **6** | **18** | **6** | **16** |
| **SD** | **35** | **51** | **1** | **2** | **1** | **5** |
| Control group |  |  |  |  |  |  |
| **Sex** | **Age (days)** | **walk (m)** | **T -l mm** | **T-t min** | **L-l mm** | **L-t min** |
| male | 50 | 0 | 7 | 14 | 9 | 9 |
| male | 51 | 0 | 6 | 20 | 7 | 15 |
| male | 67 | 0 | 7 | 14 | 5 | 12 |
| **mean** | **56** | **0** | **7** | **16** | **7** | **12** |
| **SD** | **10** | **0** | **1** | **3** | **2** | **3** |
| female | 146 | 0 | 6 | 20 | 8 | 16 |
| female | 98 | 0 | 7 | 10 | 7 | 10 |
| female | 78 | 0 | 6 | 17 | 6 | 12 |
| **mean** | **107** | **0** | **6** | **16** | **7** | **13** |
| **SD** | **35** | **0** | **1** | **5** | **1** | **3** |

### Tissue harvesting

All mice were anesthetized using isoflurane at 2-3% delivered in 95% oxygen and subjected to laminectomy of the low-thoracic and lumbar spinal segments before transcardial perfusion using ice-cold 0.1 M phosphate buffered saline (pH 7.4). Spinal cord samples were blocked into sections based on vertebral levels from thoracic (T) 6^th^ to 10^th^ levels, or from the T11^th^ to the lumbar (L) third (L2-3) vertebral levels based on evidence that these regions represent low thoracic and lumbar segments, respectively (Harrison et al., 2013). The segments were identified by counting the ribs prior to dissection. These blocks (5-8 mm long) were frozen on dry ice (see details in Table 1) and stored in at -80 Celsius.

### Western immunoblotting procedure

The spinal cord blocks were homogenized in low-binding 1.5 mL microcentrifuge tubes using a pre-sterilized Dounce glass homogenizer and ice-cold HNTG lysis buffer (50mM HEPES, 150mM NaCl, 10% glycerol, 1% Triton X-100, protease inhibitor, pH 7.5). The homogenate was first centrifuged at 1000 rpm for 10 min at 4 Celsius. The pellet (P1) was discarded, and the supernatant (S1) was removed to a new clean microcentrifuge tube and centrifuged at 14800 rpm for 45 min at 4°C. The supernatant of this fraction (S2) was carefully removed and stored in a sterile, cold Eppendorf tube. The pellet of this fraction (P2) was resuspended in the lysis buffer. The S2 fraction represented the cytosolic fraction, while the P2 fraction represented the light-membrane fraction, and both fractions were used for immunoblotting. Protein concentrations were determined using the Bradford assay (Bio-Rad, Hercules, CA).

Protein lysates were mixed with 4X Laemmli buffer (Bio-Rad, Hercules, CA) and separated using 10% Bio-Rad 10% TGX Stain-Free gel. Equal amounts of protein (24 µg) were loaded in each well. Precision Plus Protein All Blue standard (Bio-Rad, Hercules, CA) was used as markers to determine the relative molecular weights of antibody-labeled protein bands in western blots. Stain-free gels were activated for 45 seconds and imaged using a Bio-rad ChemiDoc MP imager before being transferred onto a low-fluorescence PVDF membrane using the Transblot Turbo transfer system for 7 min. Post-transferred blots were imaged using the ChemiDoc.

All blots were blocked in 5% fat-free milk or 1% BSA prepared in Tris-buffered saline, with 0.1% Tween-20 for one to two hours at room temperature. After blocking, the membranes were incubated with primary antibody: anti-CPEB3 (Abcam, cat. no. ab10883, rabbit, 1:500) or anti-anti-early endosome marker or EEA1 (Abcam, cat. no. ab2900, rabbit, 1:1000) or Na-K-ATPase (Millipore, cat. no. 06520, rabbit, 1:500) or STAT5b (Santa Cruz Biotechnology, Catalogue # SC-1656 1:300) overnight at 4°C. The membranes were washed with 1X TBST and incubated with a goat anti-rabbit HRP-conjugate secondary antibody (Abcam, cat.no. ab205718) for chemiluminescence (1 hour at room temperature). The membranes were washed and then imaged using electrochemiluminescence. The blots were incubated with anti-CPEB3, then Stat5b or Na-K-ATPase or EEA1 antibodies and stripped and washed before blocked and re-incubated with target antibody in the respective sequence. The anti-EEA1 and anti-Na/K-ATPase antibodies served as the control for checking the differentiation obtained by the subcellular fractionation.

All protein expression levels were normalized to total protein per lane using ImageLab software (Bio-Rad, Hercules, CA). Data were exported to Excel for additional normalization of combined analyses, and results are reported as mean ± standard deviation. The data in the walking and control groups were compared by Student’s t-test, after testing for normality by the Shapiro-Wilk test and by Wilcoxon-Signed Rank test when normality failed, using JAMOVI (version 2.6.) software. All original images are shown in Supplemental-Figures 1-8.

### Immunohistochemistry

Spinal cord segments were embedded in Tissue-Tek® OCT compound (Sakura Finetek) and sectioned (20 µm thick) on a cryostat (NX50, Thermo-Fisher Scientific). For immunofluorescence, the frozen slides were pre-warmed at room temperature, air-dried. Slides were incubated with 10 mM sodium citrate solution at 60-70 Celsius for 20 min and washed with 1X TBS, then permeabilized with 1.5 TBST buffer (Triton X-100, NaCl, and Trizma ® base, pH 7.6). The sections were blocked with 1% bovine serum albumin (BSA) and 10% normal donkey serum (NDS) diluted in 1.5 TBST) for an hour. Sections were incubated for one overnight at 4°C in primary antibody diluted in the blocking solution.

The following dilutions of primary antibodies were used: anti-CPEB3 (Abcam, ab10883, rabbit, 1:500); anti-NeuN (Abcam, ab104224, mouse, 1:1000; and Abcam 177487, rabbit, 1:2000) and anti-ChAT (EMD-Millipore-Sigma, AB144P, goat, 1:1000) and anti-peripherin (Chemicon-Millipore AB9282, chicken, 1:500). Then, sections were washed (3 times, 20 min in 1.5 TBST) and appropriate secondary antibody dilutions were applied at room temperature for one hour. The secondary antibodies included: Cy3-conjugated donkey anti-rabbit IgG; AlexaFlour 488-conjugated donkey anti-mouse IgG (both 1:500, from Jackson Immuno Research Laboratories, West Grove, PA, USA and ThermoFisher Scientific); or AlexaFlour488-conjugated donkey anti-rabbit (1:500, ThermoFisher Scientific, A-21206) and Cy5-conjugated donkey anti-chicken (1:500, from Jackson Immuno Research Laboratories, West Grove, PA, USA). After incubation, slides were washed and covered with Fluoromount-G with or without DAPI (Southern Biotech., Birmingham, AB, USA) to mount the slides.

Immunofluorescence was examined by a wide-field Zeiss Imager Z2 microscope using ZEN Pro image capture and analysis software (Carl Zeiss Canada, Toronto, Ontario, Canada). Data were collected as single images using 10x, 20x and 40x objective lenses. Cell counts were done using ImageJ software. Final images were assembled using CorelDraw Graphics (Corel Corp., Canada). Controls incubated with antibody diluent, without the primary antibody included) and then incubated by secondary antibodies showed no staining (Supplemental-Figure 9).

**Figure 1.**
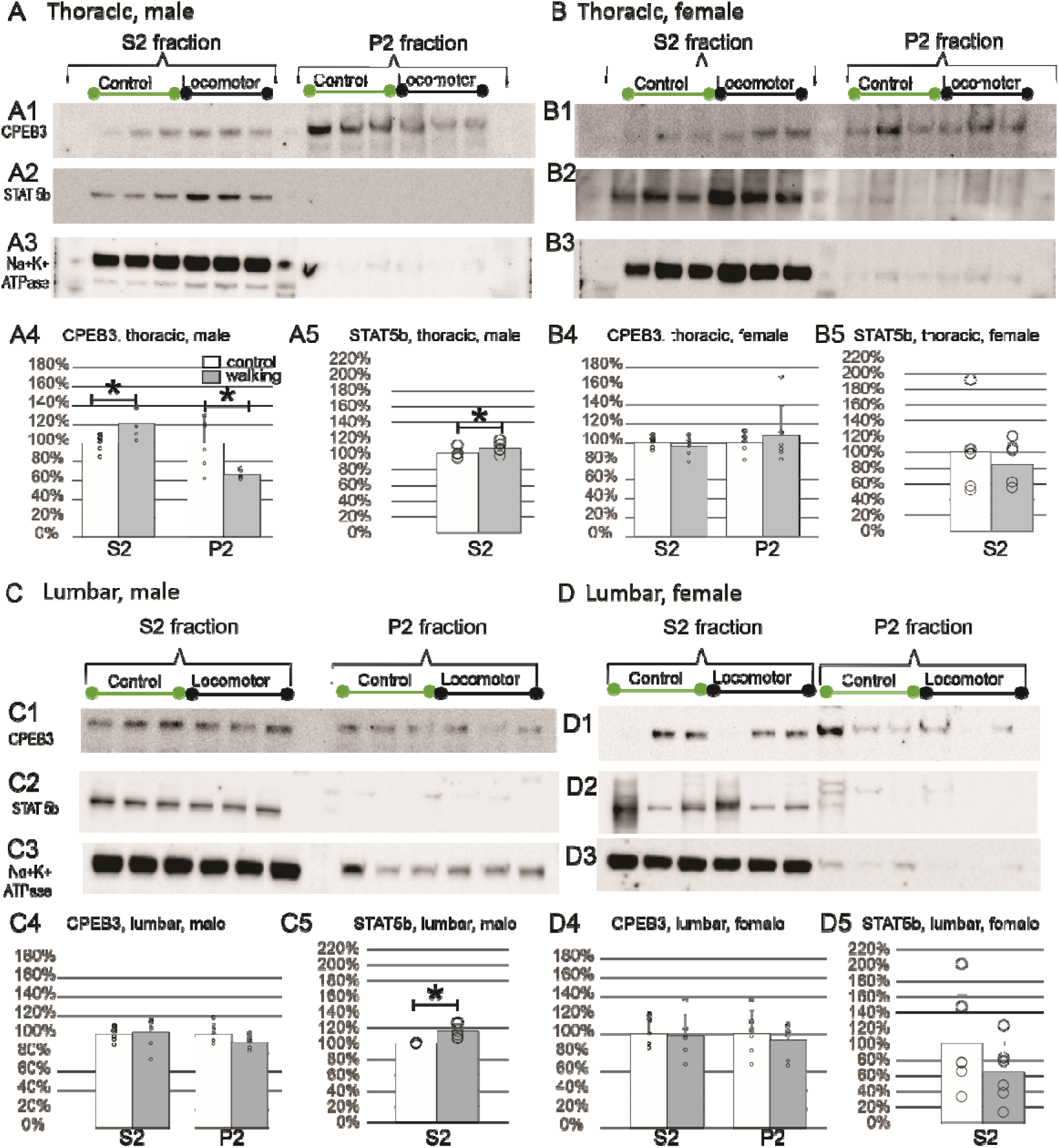
CPEB3 and STAT5B protein in thoracic and lumbar spinal cord homogenates is similar in male and female mice. The CPEB3 antibody labelling showed the presence of this protein in thoracic cord and total protein samples of all males (A, C) and females (B, D) mice (one animal per lane). The CPEB3 target (A1, B1, C1, D1) was expected at 78 kDa, the STAT5B (A2, B2, C2, D2) at 92 kDa and the NaK-ATPase (A3, B3, C3, D3) at 113 kDa in samples of soluble (S2, or cytosolic) fractions (lanes 1-6) and in the insoluble, P2, or membrane) fractions (lanes 7-12). A 10-250kD protein standard ladder was used as a reference of size and loading per lane was 24 µg. The bar graphs show quantitative, normalized data expressed as % control (i.e. levels in non-locomotor mice) for each comparison of S2 or P2 fraction, in males or females in thoracic or lumbar samples, as indicated in each panel. Normalization of target protein to total protein (stain-free blot) was performed for CPEB3 (A4, B4, C4 and D4) and for STAT5B (A5, B5, C5, D5). Circles represent individual data points, error bars indicate SD; star indicates significant differences (p<0.05).

### CPEB3 protein expressed in thoracic and lumbar spinal cord

This study examined the presence of CPEB3 protein in thoraco-lumbar spinal segments. To determine whether CPEB3 was cytosolic or present in light-membrane fractions, separation of the membrane and cytosolic fractions after whole-tissue homogenization was verified. To examine changes in CPEB3 levels after neural activity, we compared semi-quantitative protein expression in age-matched pairs of C57BL/6 wild-type mice after 120 minutes of walking on a treadmill, on average 741±66 m, 90 minutes prior to tissue collection. The thoracic and lumbar cord data from independent blots showed similar results with respect to behavior i.e. resting or walking for 2 hrs. Results in Fig. 1 show that the CPEB3 protein (Fig. 1A1, 1B1, 1C1 and 1D1) was detected in both the soluble fractions (S2) and in the insoluble fractions (P2) similarly in males (Fig. 1A, 1C) and females (Fig. 1B, 1D) in samples collected from the thoracic (Fig. 1A-1B) and from the lumbar segments (Fig. 1C-1D). The STAT5B target band (Fig. 1A2, 1B2, 1C2, 1D2) was present in the cytosolic (S2) fraction in both thoracic and lumbar samples. The presence of sodium-potassium-ATPase (Na/K/ATPase) mainly in the soluble (S2) fractions was used as a verification that fractionation segregated proteins as reported earlier (Schimmel et al., 1973).

Semi-quantification of CPEB3 expression in tissue from mice subjected to walking compared with control (non-locomoting) mice showed that CPEB3 increased in soluble fractions (S2, Fig. 1A4) by 20.7±15% (t-test, p=0.03, Table 3). Comparison of STAT5b showed that in males, in the thoracic segments, there was a significant increase in mice that were subjected to walking, by 6.8±0.1% compared to levels in the non-walking mice (t-test, p=0.023). In females, however, there was no evidence for significant changes in thoracic CPEB3 or STAT5b (Fig. 1B4, 1B5, see also Table 3). From samples originating from the lumbar spinal cord, there was a similar trend in males to the pattern in thoracic cord but without reaching a significant level of change (Table 2). STAT5b in the lumbar soluble fraction showed increased significantly by 15.7±0.1% (paired t-test, p=0.01). In lumbar cords of females, there were no significant changes in either CPEB3 (Fig. 1D4) or in Stat5B (Fig. 1D5) levels (see also Tables 2 and 3). Comparison of CPEB3 in the insoluble fractions showed no significant change in any of the tested segments. Stat5b was not detected at all in the insoluble fractions under these conditions.

**Table 2.** Summary of statistics examining CPEB3. Descriptive statistics and comparison of results obtained from soluble (s2), or insoluble (p2) samples extracted from control (ctrl) or locomotor (loco) mice. Abbreviations: T-thoracic, L-lumbar, m-male, f-female, N=tissue samples, SD-standard deviation, SE-standard error, p-the p value at alpha=0.05 of Student’s t-test. Asterisks indicate statistically significant differences (p<0.05).

| CPEB3 | sex | sample | N | Mean | Median | SD | SE | p |
|---|---|---|---|---|---|---|---|---|
| T | m | s2-ctrl | 6 | 1 | 1.03 | 0.0901 | 0.0368 | **0.034**\* |
|  |  | s2-loco | 6 | 1.207 | 1.175 | 0.149 | 0.0608 |  |
|  |  | p2-ctrl | 6 | 1.002 | 1.055 | 0.2684 | 0.1096 | **0.039**\* |
|  |  | p2-loco | 6 | 0.662 | 0.645 | 0.0475 | 0.0194 |  |
|  | f | s2-ctrl | 6 | 1 | 1.01 | 0.066 | 0.027 | 0.462 |
|  |  | s2-loco | 6 | 0.957 | 0.975 | 0.1076 | 0.0439 |  |
|  |  | p2-ctrl | 6 | 1 | 1.03 | 0.1121 | 0.0458 | 0.692 |
|  |  | p2-loco | 6 | 1.07 | 0.95 | 0.3234 | 0.132 |  |
| L | m | s2-ctrl | 6 | 0.958 | 0.97 | 0.1532 | 0.0625 | 0.231 |
|  |  | s2-loco | 6 | 1.018 | 1.1 | 0.1734 | 0.0708 |  |
|  |  | p2-ctrl | 6 | 1.002 | 0.97 | 0.1089 | 0.0445 | 0.123 |
|  |  | p2-loco | 6 | 0.913 | 0.92 | 0.07 | 0.0286 |  |
|  | f | s2-ctrl | 6 | 1.002 | 0.95 | 0.152 | 0.062 | 0.849 |
|  |  | s2-loco | 6 | 0.975 | 0.96 | 0.2358 | 0.0963 |  |
|  |  | p2-ctrl | 6 | 1.002 | 1.005 | 0.2451 | 0.1 | 0.387 |
|  |  | p2-loco | 6 | 0.932 | 1.015 | 0.1953 | 0.0797 |  |

**Table 3.** Summary of statistics examining Stat5b. Descriptive statistics and comparison of results obtained from soluble (s2) samples extracted from control (ctrl) or locomotor (loco) mice. Abbreviations: T-thoracic, L-lumbar, m-male, f-female, N=tissue samples, SD-standard deviation, SE-standard error, p-the p value at alpha=0.05 of Student’s t-test. Asterisks indicate statistically significant differences (p<0.05).

| STAT5B | sex | sample | N | Mean | Median | SD | SE | p |
|---|---|---|---|---|---|---|---|---|
| T | m | s2-ctrl | 6 | 1 | 1 | 0.06261 | 0.02556 | **0.023**\* |
|  |  | s2-loco | 6 | 1.068 | 1.05 | 0.0725 | 0.0296 |  |
|  | f | s2-ctrl | 6 | 1.002 | 0.985 | 0.50261 | 0.20519 | 0.527 |
|  |  | s2-loco | 6 | 0.845 | 0.82 | 0.28077 | 0.11462 |  |
| L | m | s2-ctrl | 6 | 0.997 | 0.995 | 0.00816 | 0.00333 | **0.007**\* |
|  |  | s2-loco | 6 | 1.157 | 1.13 | 0.09288 | 0.03792 |  |
|  | f | s2-ctrl | 6 | 1 | 0.76 | 0.62568 | 0.25543 | 0.296 |
|  |  | s2-loco | 6 | 0.638 | 0.63 | 0.39509 | 0.1613 |  |

**Figure 2.**
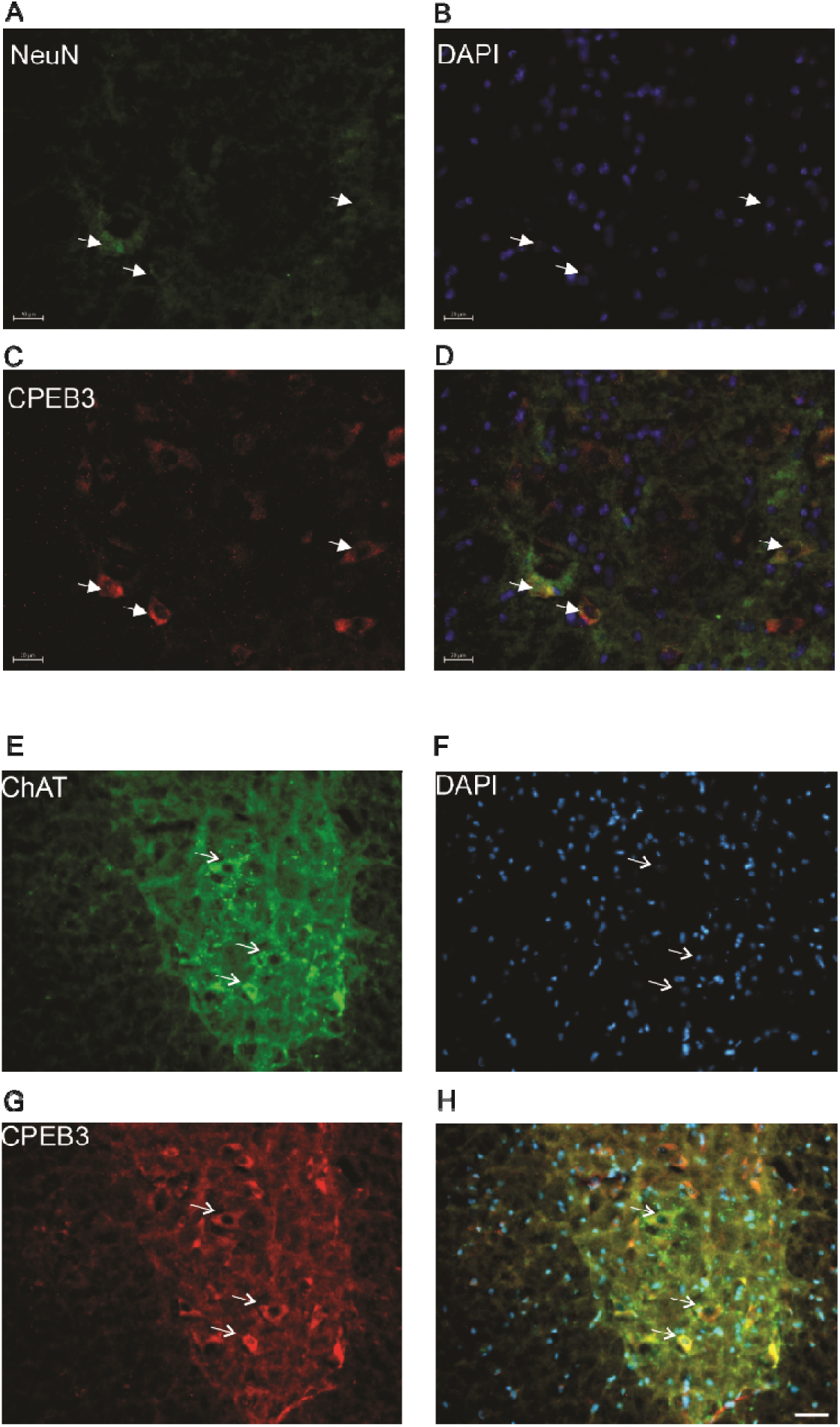
Expression of CPEB3 in neurons of the ventral horn. Immunofluorescence labelling of thin lumbar spinal cord sections with antibodies to detect NeuN green, in A and CPEB3, red in B, and the nuclear DAPI, blue in C, with overlay of all colors shown in D. From another mouse, immunofluorescence labelling of thin lumbar spinal cord sections with antibodies to detect choline acetyltransferase, ChAT, green in E and CPEB3, red in F, and DAPI, blue in G with overlay of all colors shown in H. Scale bars represent 20 µm in A-D and 50 µm in F-H and arrows indicate neurons positive for CPEB3 and NeuN in A-D, and CPEB3 and ChAT in E-H.

### Cholinergic neurons of the ventral horn express CPEB3 protein

Cytosolic expression of CPEB3 was confirmed by immunofluorescence labeling in thin, transverse sections of mouse spinal cord. The cells positive for CPEB3 were often labelled for NeuN (Fig. 2). This was similar along the entire rostro-caudal length of both thoracic and lumbar segments examined. CPEB3 protein strongly expressed in cells within the ventral horn. NeuN-positive cells, including large ventral horn neurons were the most prominently labeled structures (Fig. 2E). After quantifying the number of NeuN-positive cells from serial sections, 88±5% (of 304 cells total) was found to be positive for CPEB3.

The appearance of CPEB3 associated with cells positive for choline acetyltransferase, ChAT, is shown in Fig. 2 in a section labeled for both CPEB3 and ChAT. CPEB3 was frequently localized to ChAT-positive somas and the immunofluorescence labeling for CPEB3 in mouse spinal cord transverse sections was simultaneously labeled for NeuN. This was similar along the entire rostro-caudal length of both low-thoracic and lumbar segments examined. After quantifying from serial sections, the number of NeuN positive cells, 88±5% were also positive for CPEB3 (n=304 cells total). No major differences appeared in the density or distribution of CPEB3 associated with these cell groups at different thoracic spinal levels, although this was not examined quantitatively. The NeuN-positive somas of large ventral horn neurons with dendrites reaching into the white matter were suggestive of motoneurons expressing CPEB3. We, therefore examined this using an antibody against peripherin. These experiments showed that the large ChAT+ neurons shown in Fig. 3, which were positive for CPEB3, also expressed peripherin. This was interpreted as an indication of ventral spinal motoneurons with peripherally-projecting axons express CPEB3.

**Figure 3.**
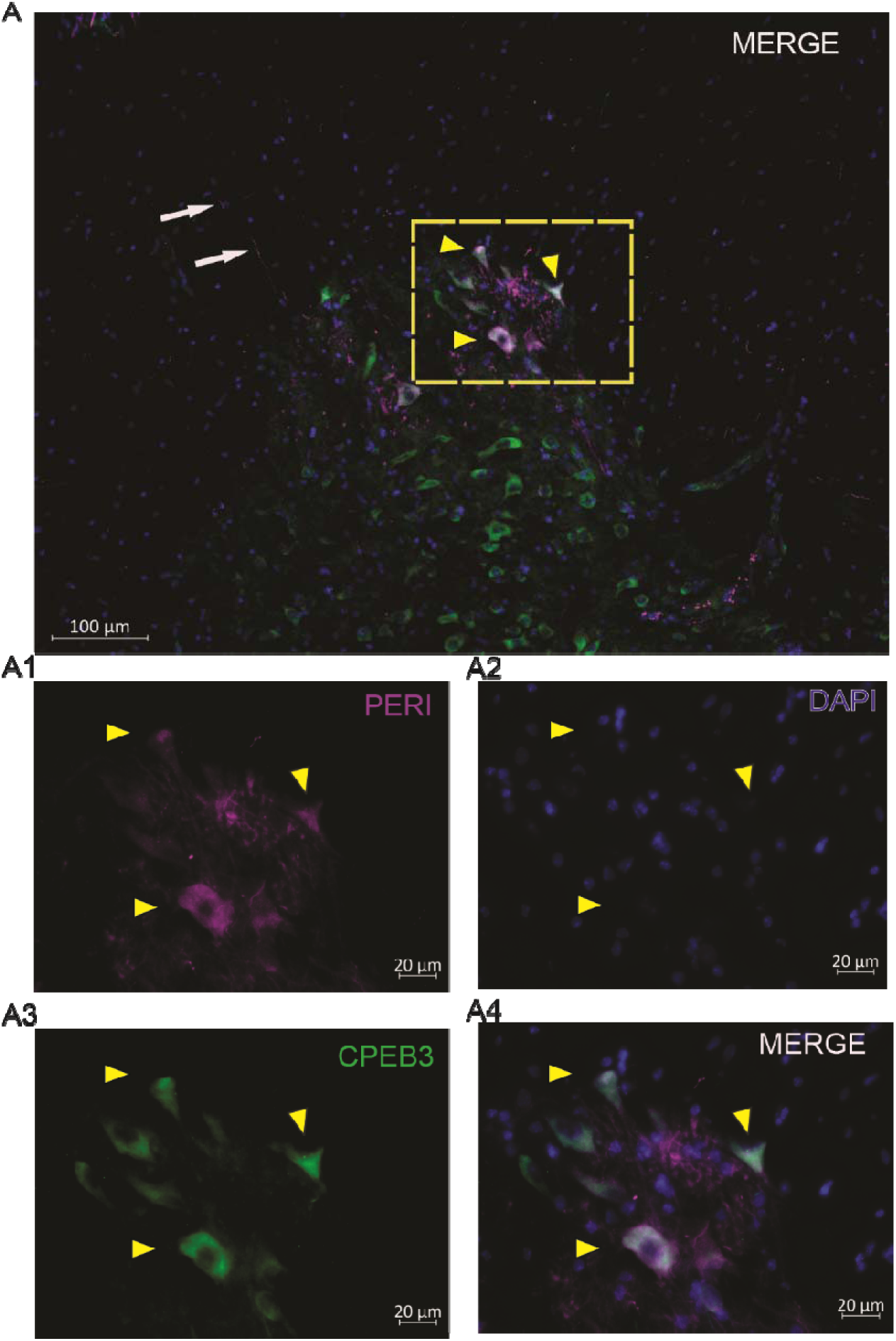
The presence of CPEB3 in adult mouse motoneurons. Immunofluorescence microscopy images from the spinal grey matter showing the ventral horn (A) and the boxed region imaged at higher magnification (A1-A4). The axons (arrows) and somas (arrowheads) of some cells labelled with antibodies to peripherin (A1, magenta, PERI); The CPEB3 (green, A3) and nuclear DAPI (blue, A2) shown overlayed on the merged panel (A4) illustrating th colocalization of these proteins. Scale bar in A is 100 µm, in A1-A4 is 20 µm.

## Discussion

The presence of CPEB3 in dorsal root ganglia at cervical, thoracic, and lumbar levels has been previously established (Fong et al., 2016). This study found CPEB3 protein expression in adult mouse spinal grey matter in soluble and insoluble fractions of spinal cord segments ranging from the low-thoracic (T6^th^) to mid-lumbar (L3^rd^) segments. Additional immunohistochemical tests on thin spinal cord sections substantiated these findings, showing that in the ventral horn, CPEB3 is expressed in neurons containing ChAT, which is indicative of a cholinergic phenotype. Moreover, spinal motoneurons positive for peripherin, an intermediate filament protein indicative of axons reaching the peripheral nervous system, were also labeled positive for CPEB3.

To our knowledge, this study is the first to examine activity-induced changes in spinal CPEB3 levels after walking. The neural activity induced by two hours of low-speed treadmill walking (8-15 cm/s) increased CPEB3 soluble fractions of the thoracic spinal segments in male mice, but it did not significantly alter CPEB3 expression in female mice. Here, we examined only one time point (120 minutes of walking) based on previously recognized upregulation of c-Fos early gene expression in response to walking (Ahn et al., 2006; Barth et al., 2004; Dai et al., 2005). However, we cannot rule out the possibility that CPEB3 levels change when mice are required to perform longer-duration or higher-intensity locomotion. Indeed, examining CPEB3 levels at various time points relative to the onset of a motor task would be necessary. The levels we tested here correspond to low-level walking activity (Lemieux et al., 2016) and higher walking speeds would be expected to recruit additional cholinergic interneurons (Zagoraiou et al., 2009). CPEB3 expression was similar in males and females and our data suggests potential sex-related differences, although, our sample size was small, and validation of sex-based differences is necessary with more samples. Moreover, the age may also be critical to control for, as one report showed increased spinal c-Fos immunoreactivity in aged mice and dogs (Ahn et al., 2015).

Examination of oocyte extracts showed that most CPEB proteins are associated with various membrane-spanning proteins (Cao et al., 2005). Localization of CPEB3 separately in soluble (S2) and insoluble (P2) fractions obtained from homogenized spinal cord tissue was expected based on what is known about CPEB3’s subcellular localization. This finding is also consistent with data from cultured neurons (Ford et al., 2019) showing that within the cytoplasm, basal CPEB3 in neurons resided in fractions containing the P-body component. P-bodies in neurons are thought to be cytoplasmic granules that specialize in both translational silencing and mRNA degradation (Zeitelhofer et al., 2008). One hypothesized role of the P-bodies is to sequester mRNAs and keep them translationally silenced. Dendritic P-body-like structures are present in the soma and dendrites of mammalian neurons and exhibit re-localization along the dendrites to distant sites in response to synaptic activation (Cougot et al., 2008). P-bodies may regulate local translation (Ford et al., 2019). The idea that CPEB3 translocates to polysomes after spinal neural activation needs to be further explored using specific markers, as our results imply this possibility given that CPEB3 was found in the insoluble (membrane) fractions.

Less information is available on the locomotor-activity-dependent modulation of STAT5. Neuronal development, however, is greatly modulated by this protein. For example, phosphorylated STAT5 (pSTAT5) has been described to be present in telencephalic neurons, and axons and its absence leads to fewer of interneurons in the cortical marginal zone and misguided axonal development in the brain (Markham et al., 2007). Moreover, STAT5a/b are expressed in commissural neurons of the embryonic spinal cord and likely play a key role as a guidance molecule. Since the nuclear translocation of CPEB3 is linked to STAT5b as a perceived switch for downstream signaling pathway regulation (Peng et al., 2010) we probed STAT5b in spinal cord tissue. We found a clear increase in males related to locomotor activity. Understanding the conformational dynamics and molecular mechanisms governing the CPEB3–STAT5b complex is therefore essential to uncovering how nuclear-cytoplasmic signaling coordinates neuronal function and how synapse guidance and stabilization occur in the spinal commissural system.

Given the spinal cord’s involvement in motor behavior and it’ role in transmitting sensory information to the brain, spinal cord plasticity is thought to connect more directly to activity and behavior than elsewhere in the central nervous system (Wolpaw, 2010). Key questions about motor memory as defined by Wolpaw (2010) include 1) how protein synthesis/degradation is regulated in molecular terms and 2) which spinal neural populations are equipped with machinery that is unique in terms of plasticity-related proteins. This study has demonstrated that CPEB3 and linked machinery may be a candidate protein for steering motor activity-driven plasticity. Once the cytoplasmic polyadenylation machinery is activated, it triggers phosphorylation, polyadenylation, and translation of several cytoplasmic polyadenylation-containing mRNAs. Future studies need to address STAT5 - CPEB3 interactions in both classical and non-conventional pathways to better understand the formation and retention of spinal synaptic connectivity.

## Conclusion

This study demonstrated for the first time that CPEB3 is expressed in cholinergic thoracic and lumbar spinal interneurons, including cholinergic motoneurons of mature mice. Two hours of treadmill walking was sufficient to alter relative CPEB3 and STAT5b protein levels in neurons of the tested thoraco-lumbar segments; however, these findings warrant further research into CPEB3-controlled translation relates to motor activity-associated plasticity in spinal neural networks. Examining in more detail any sex-based differences and which mechanisms stabilize synapses after locomotor training could be examined by building on our findings of CPEB3 expression in spinal cholinergic cells.

## Author Contributions

The contributions of individual authors referred to by their initials were as follows. The study was conceived and designed by K.S.; locomotor activity tasking and tissue harvesting was performed by P. L., S.B., P.S. under the supervision of X.C. and K.S. The immunohistochemical protocols were developed by S.B. and P.L. under the supervision of X. C. and K.S. and the blotting protocols were developed by S.B. and P.S. under the supervision of K.S. Cell counting was performed by F.S.; S.B. and P.S. Figures, tables and the manuscript draft was prepared by P. L., P.S. and K.S. and all authors contributed to the submitted version. All authors agree to be accountable for the content of this work.

## Acknowledgements

The Authors would like to thank Drs. James I. Nagy and Robin Da Silva for help with microscopy and equipment and resources sharing, Dr. Bruce Lynn for occasional technical help.

## Ethics statement

All experiments were approved by the University of Manitoba Central Animal Care Committee prior to commencing the experiments in accordance with the ethical standards set by the Canadian Council on Animal Care.

## Data availability statement

The original data can be accessed at the following link: https://borealisdata.ca/dataverse/CPEB3

