## Supplemental-Figures1-9 for "Cytoplasmic polyadenylation protein-3 is expressed in adult mouse cholinergic spinal neurons, including motoneurons"

A
Total protein,
Fig. 1A1_2

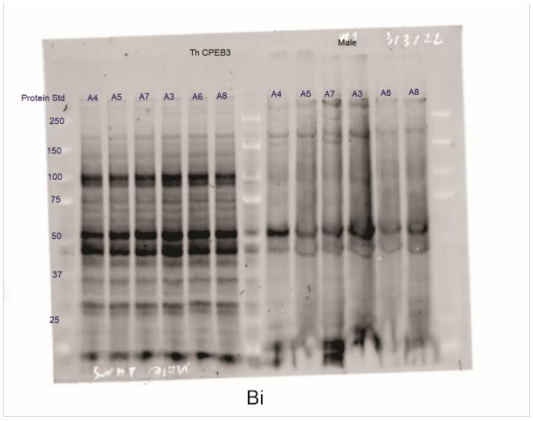


B
Fig. 1, panel A1

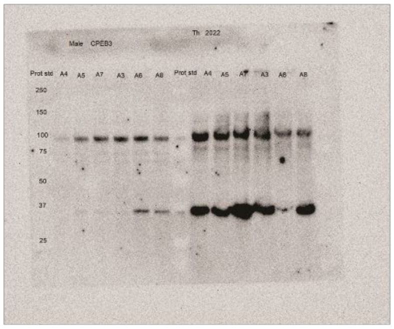


Bi

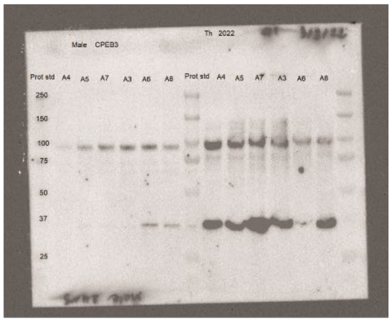


C
Fig. 1, panel A2

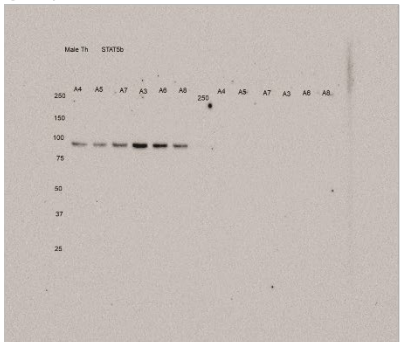


Ci

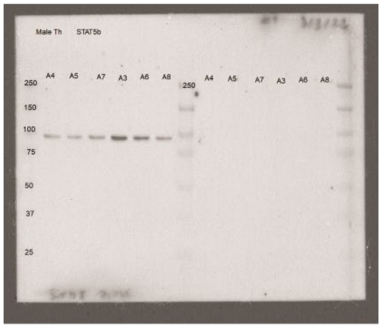


**Supplemental-Figure 1. Images of original and annotated western blot results.** Total protein corresponding to results shown in Figure 1A1 and 1A2 in A, results with the CPEB3 in B and with the Stat5b antibody in C. Note that S2, or cytosolic fractions were in lanes 1-6 and the insoluble, P2, or membrane fractions were in lanes 7-12. A 10-250 kD protein standard ladder was used as a reference of size and loading per lane was 24 µg.

A
Total protein
Fig. 1B1_2

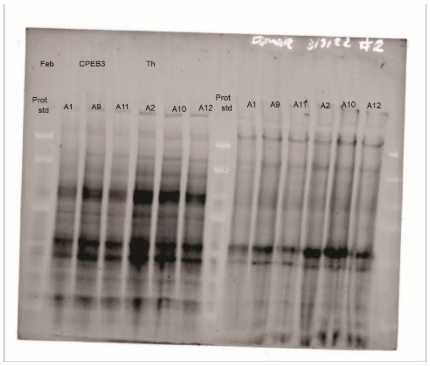


B
Fig. 1, panel B1

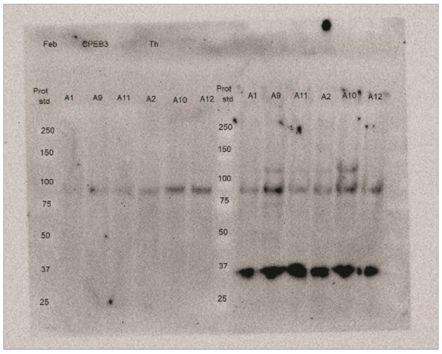


Bi

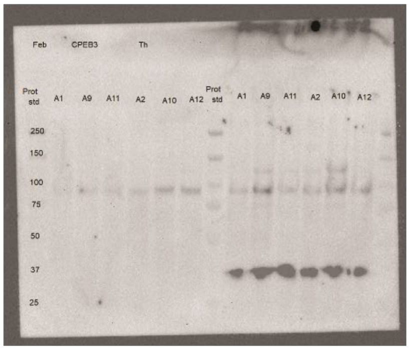


C
Fig. 1, panel B2

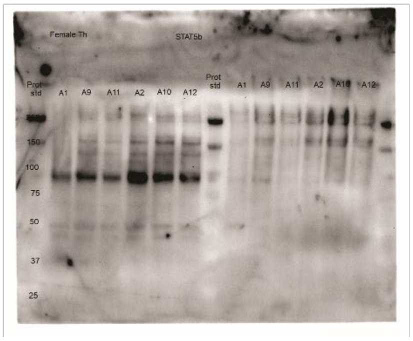


Ci

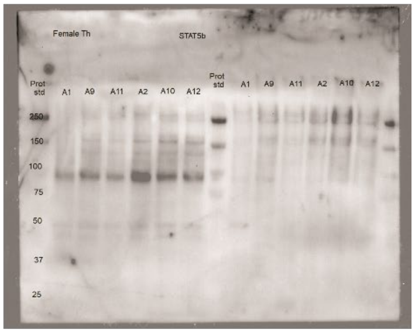


**Supplemental-Figure 2. Images of original and annotated western blot results.** Total protein corresponding to results shown in Figure 1B1 and 1B2 in A, results with the CPEB3 in B and with the Stat5b antibody in C. Note that S2, or cytosolic fractions were in lanes 1-6 and the insoluble, P2, or membrane fractions were in lanes 7-12. A 10-250 kD protein standard ladder was used as a reference of size and loading per lane was 24 µg.

A
Total protein, Fig. 1C1_2

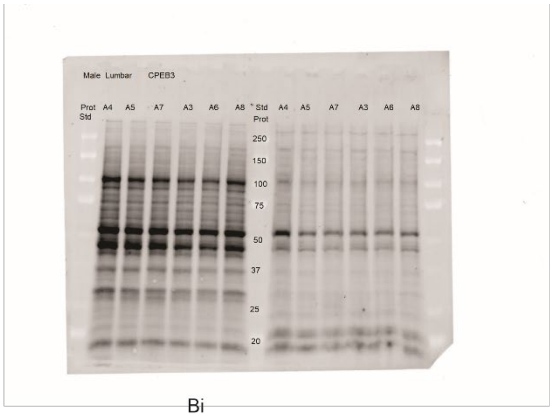


B
Fig. 1, panel C1

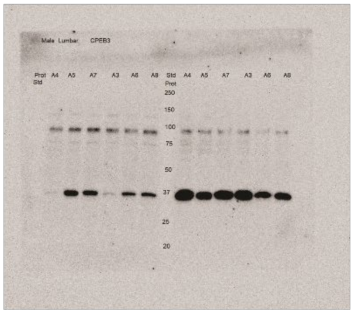


Bi

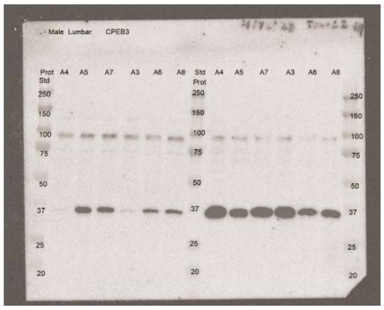


C
Fig. 1, panel C2

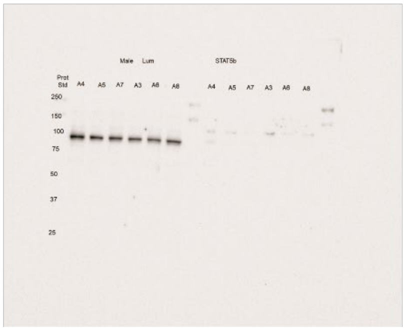


Ci

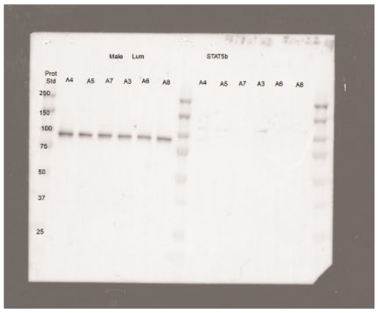


**Supplemental-Figure 3. Images of original and annotated western blot results.** Total protein corresponding to results shown in Figure 1C1 and 1C2 in A; results with the CPEB3 in B and with the Stat5b antibody in C. Note that S2, or cytosolic fractions were in lanes 1-6 and the insoluble, P2, or membrane fractions were in lanes 7-12. A 10-250 kD protein standard ladder was used as a reference of size and loading per lane was 24 µg.

A
Total protein,
Fig. 1D1_2

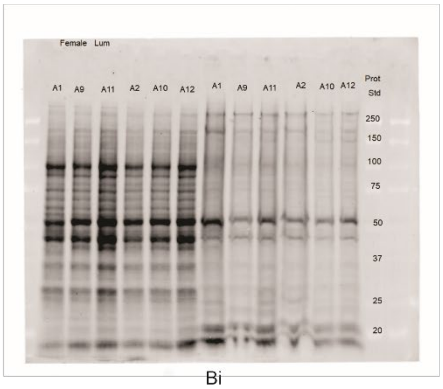


B
Fig. 1, panel D1

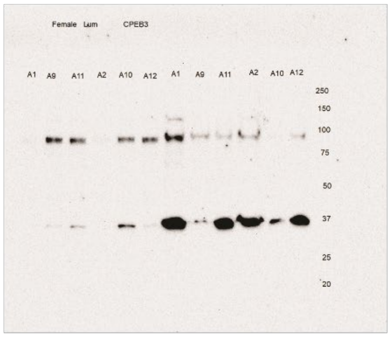


Bi

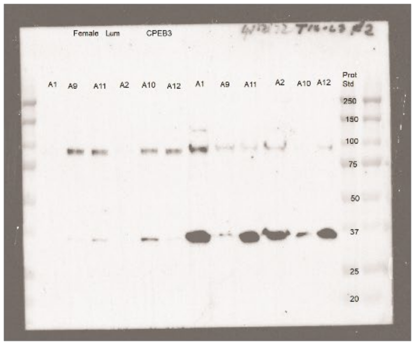


C
Fig. 1, panel D2

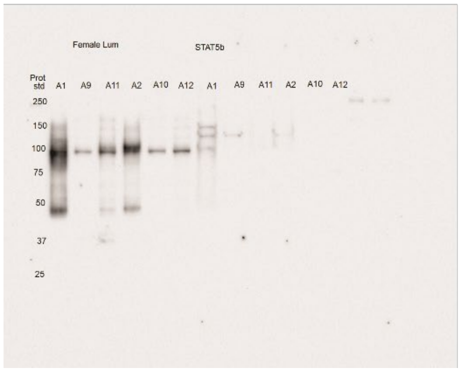


Ci

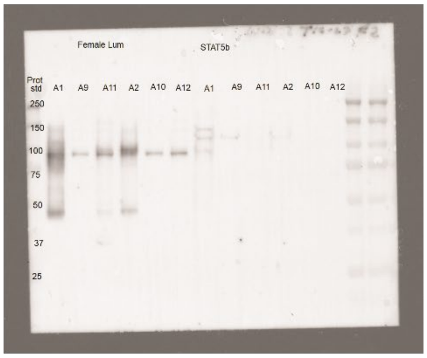


**Supplemental-Figure 4. Images of original and annotated western blot results.** Total protein corresponding to results shown in Figure 1D1 and 1D2 in A, results with the CPEB3 in B and with the Stat5b antibody in C. Note that S2, or cytosolic fractions were in lanes 1-6 and the insoluble, P2, or membrane fractions were in lanes 7-12. A 10-250 kD protein standard ladder was used as a reference of size and loading per lane was 24 µg.

A
total protein, second experiment, panels Fig. 1A4_5 for quantification

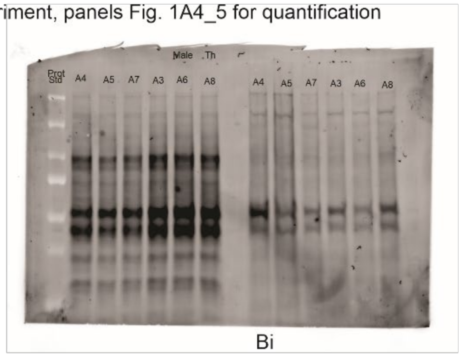


B
Fig. 1, panel A4

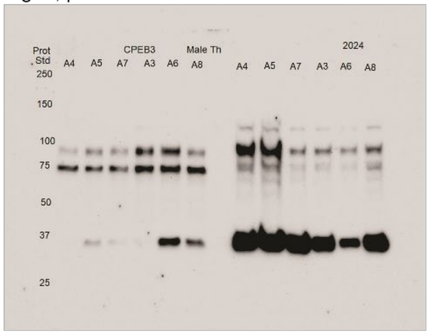


Bi

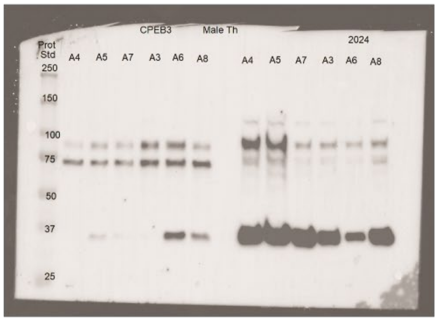


C
Fig. 1, panel A5

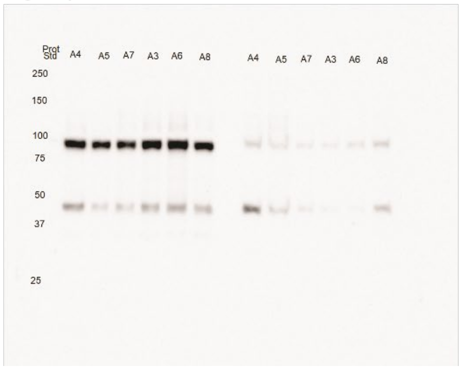


Ci

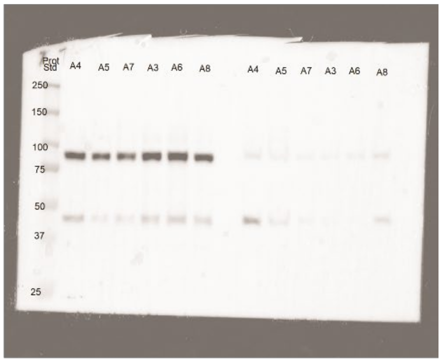


**Supplemental-Figure 5. Images of original and annotated western blot results.** Total protein corresponding to second experiment results (not shown) but used for quantification reflected in in panels of Figure 1A4 and 1A5 in A, results with the CPEB3 in B and with the Stat5b antibody in C. Note that S2, or cytosolic fractions were in lanes 1-6 and the insoluble, P2, or membrane fractions were in lanes 7-12. A 10-250 kD protein standard ladder was used as a reference of size and loading per lane was 24 µg.

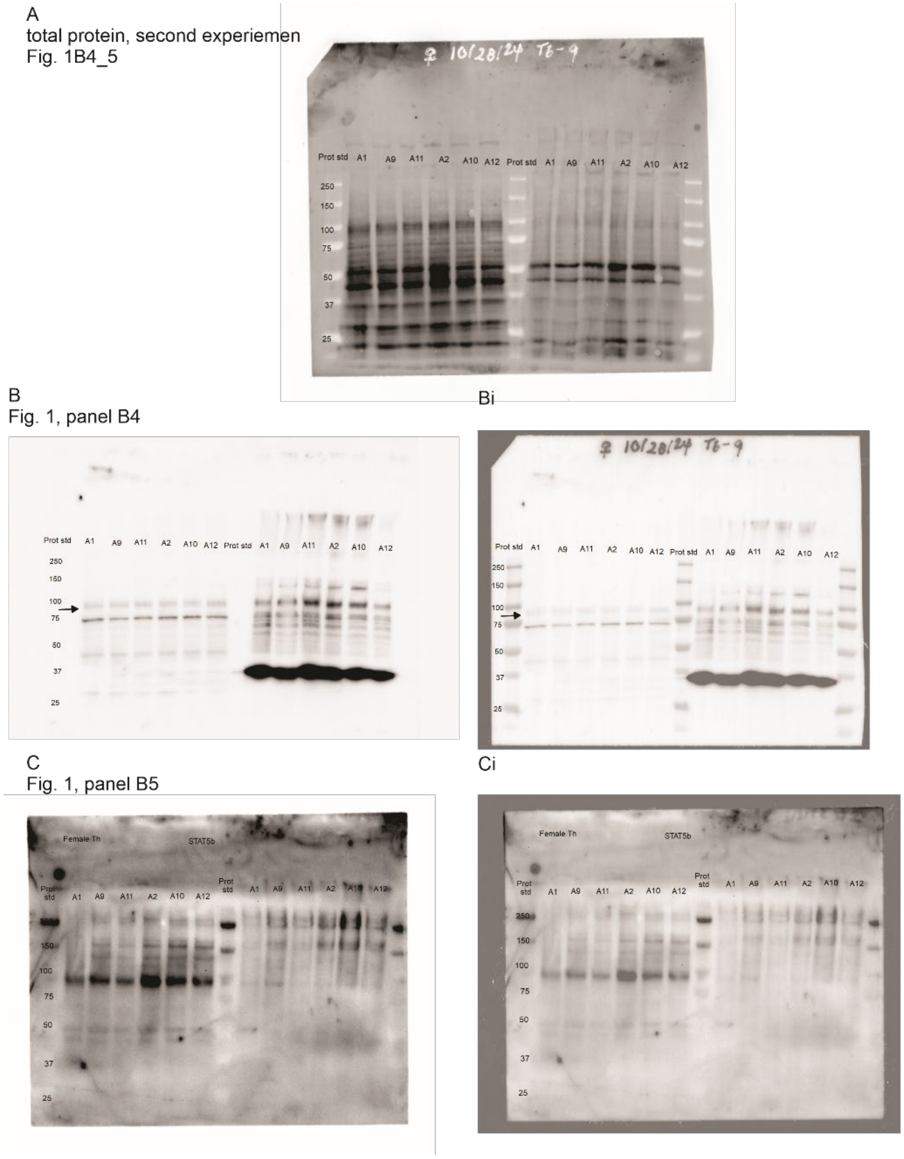


**Supplemental-Figure 6. Images of original and annotated western blot results.** Total protein corresponding to second experiment results (not shown) but used for quantification reflected in in panels of Figure 1B4 and 1B5 in A, results with the CPEB3 in B and with the Stat5b antibody in C. Note that S2, or cytosolic fractions were in lanes 1-6 and the insoluble, P2, or membrane fractions were in lanes 7-12. A 10-250 kD protein standard ladder was used as a reference of size and loading per lane was 24 µg.

A
second experiment
Total protein,
Fig. 1C4_5

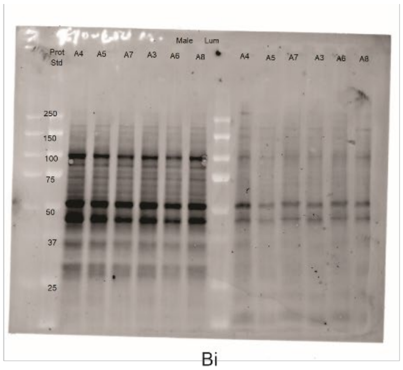


B
Fig. 1, panel C4

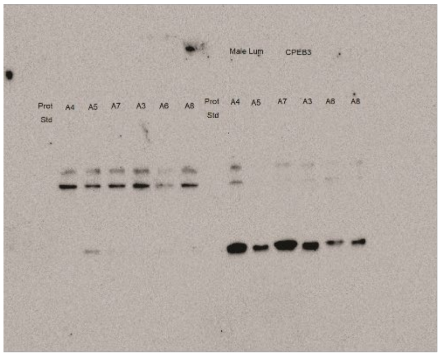


Bi

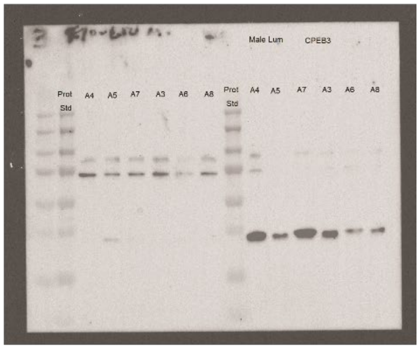


C
Fig. 1, panel C5

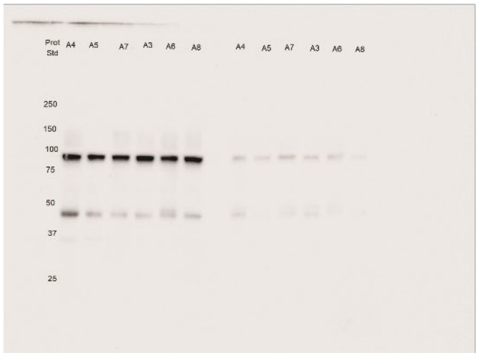


Ci

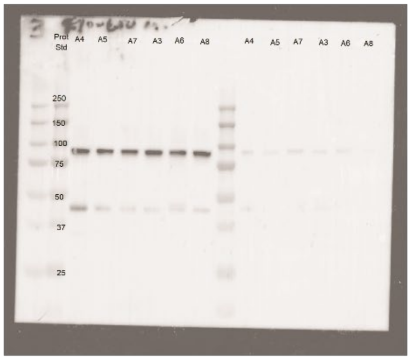


**Supplemental-Figure 7. Images of original and annotated western blot results.** Total protein corresponding to second experiment results (not shown) but used for quantification reflected in in panels of Figure 1C4 and 1C5 in A, results with the CPEB3 in B and with the Stat5b antibody in C. Note that S2, or cytosolic fractions were in lanes 1-6 and the insoluble, P2, or membrane fractions were in lanes 7-12. A 10-250 kD protein standard ladder was used as a reference of size and loading per lane was 24 µg.

A
Total protein, second experiment
Fig. 1D4_5

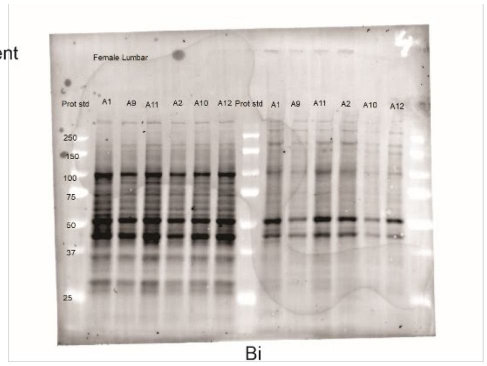


B
Fig. 1, panel D4

Bi

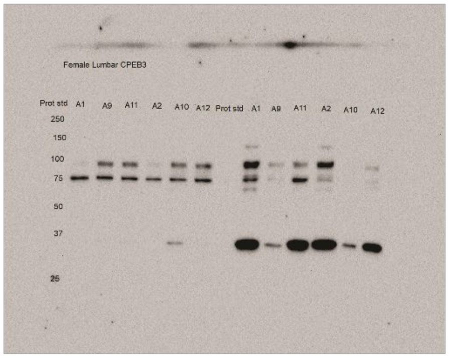


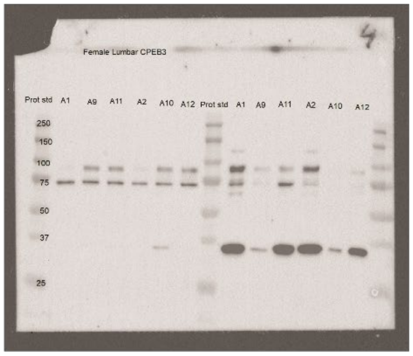


C
Fig. 1, panel D5

Ci

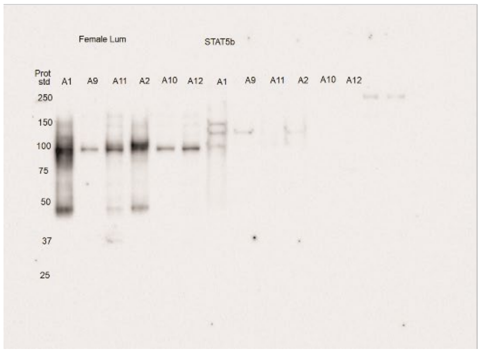


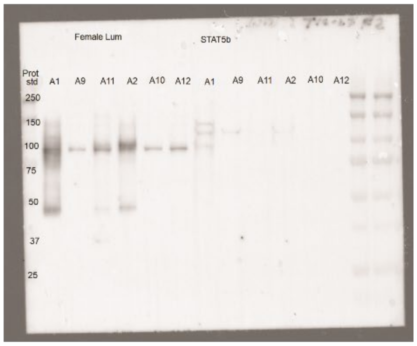


**Supplemental-Figure 8. Images of original and annotated western blot results.** Total protein corresponding to second experiment results (not shown) but used for quantification reflected in in panels of Figure 1D4 and 1D5 in A, results with the CPEB3 in B and with the Stat5b antibody in C. Note that S2, or cytosolic fractions were in lanes 1-6 and the insoluble, P2, or membrane fractions were in lanes 7-12. A 10-250 kD protein standard ladder was used as a reference of size and loading per lane was 24 µg.

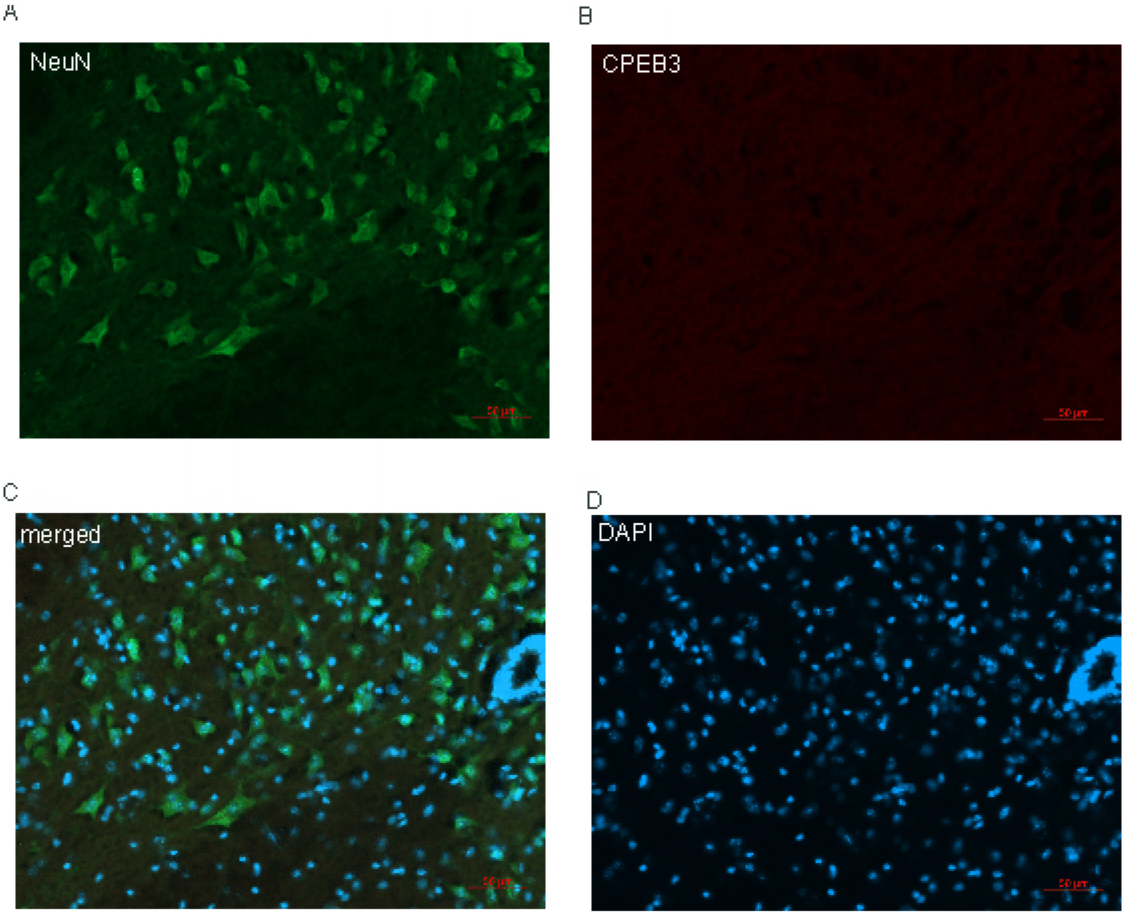


**Supplemental-Figure 9. Primary CPEB3 omission results.** Immunofluorescence microscopy images when primary CPEB3 antibodies were omitted showing the central canal and ventral horn cells labelled with antibodies to NeuN (green) in A; CPEB3 (red) in B, and nuclear DAPI (blue) in D; with the overlayed, merged channels shown in C. Scale bar is 50 µm.
